# sstar2: A Python Package for *S*^*^-based Archaic Introgression Detection with Machine Learning

**DOI:** 10.64898/2026.05.31.729079

**Authors:** Andrea Koça, Alexander Stöckl, Simon Chen, Martin Kuhlwilm, Xin Huang

## Abstract

Detecting introgressed genomic fragments from unsampled or extinct source populations remains challenging. The *S*\* statistic is widely used for this purpose, but the original sstar implementation relies on generalized additive models to smooth quantile-specific values precomputed from fixed count bins, requiring simulations with fixed numbers of segregating sites. Here, we present sstar2, a Python update that replaces this procedure with quantile regression to directly estimate *S*\* thresholds at specified null quantiles from simulated genomic windows. We benchmarked sstar2 against the original sstar, linear quantile regression, and random forest quantile regression across three demographic models with both phased and unphased simulated data. sstar2 showed the best overall performance among the evaluated methods, with the most pronounced improvement under a challenging demographic model of ghost introgression in bonobos. These results show that sstar2 improves *S*\* threshold calibration while making *S*\*-based introgression analyses more flexible and compatible with modern simulation workflows.

## 1 Introduction

Gene flow between divergent populations can leave distinct genomic fragments, but detecting such fragments is challenging when the introgressing source population is unsampled or extinct (Huang et al. 2025). The *S*\* statistic was introduced to detect introgressed fragments by identifying patterns of linked variation expected under introgression (Plagnol and Wall 2006). Since then, *S*\* has been applied to both human and non-human genomic data, but earlier implementations were limited in flexibility and usability (Vernot and Akey 2014; Vernot et al. 2016; Kuhlwilm et al. 2019; Pawar et al. 2023). The sstar package addressed this limitation by providing a Python implementation for calculating *S*\* across genomic windows and identifying candidate introgressed regions across user-defined demographic settings (Huang et al. 2022).

In this framework, *S*\* scores observed in empirical data are compared with null expectations estimated from simulations under demographic models without introgression. These null expectations are modeled using generalized additive models (GAMs), with simulated *S*\* scores as the response and window-level mutation counts, local recombination rates, and a user-specified quantile of the simulated null distribution as predictors (Vernot and Akey 2014; Vernot et al. 2016; Huang et al. 2022). To cover a range of mutation counts, sstar fits expected *S*\* scores using simulations stratified into predefined count bins, which requires generating replicate data sets with fixed numbers of segregating sites using ms (Hudson 2002). Under this conditional simulation mode, the total number of segregating sites in each simulated genomic window is fixed by the chosen bin rather than being determined by the mutation rate and sequence length.

The use of fixed count bins highlights a key limitation of this GAM-based calibration: it does not estimate conditional quantiles directly. Instead, quantile-specific expected *S*\* values are precomputed within each bin, and the GAM then performs smooth interpolation over these values. To address this limitation, we implemented sstar2 (version 2.0.0), replacing the GAM-based calibration with quantile regression to directly estimate *S*\* thresholds at specified null quantiles from simulated genomic windows. This also removes the need to construct fixed count bins using ms and enables integration with modern population genetic simulation workflows based on flexible simulators such as msprime and curated demographic models from stdpopsim (Adrion et al. 2020; Baumdicker et al. 2022; Lauterbur et al. 2023; Gower et al. 2025).

## 2 Materials and Methods

### 2.1 Overview of sstar2

sstar2 is implemented in Python and relies on msprime for simulation, scikit-allel for processing genotype data in VCF format (Danecek et al. 2011), and scikit-learn for quantile regression. The package runs on Unix/Linux operating systems and provides two commands, train and infer: train fits a gradient boosting quantile regressor to simulated data, whereas infer uses the trained model to predict *S*\* thresholds at specified null quantiles. sstar2 assumes biallelic genotype data with derived alleles coded as 1 and supports both phased and unphased data. In sstar2, the reference population denotes the population without introgressed fragments, whereas the target population denotes the population receiving introgressed fragments. Only private mutations—variants present in the target population but absent from the reference population—are used to compute *S*\* scores.

### 2.2 Input Files

For training, sstar2 requires a Demes YAML file specifying the demographic model without introgression used for simulation (Gower et al. 2022), together with a YAML configuration file. The configuration file is organized into three sections. The simulation section defines the simulation batch size, reference and target sample sizes, reference and target population names, sequence length, mutation and recombination rates, ploidy, phasing status, number of simulated genomic windows, number of processes, and random seed. The preprocessing section specifies the VCF file, chromosome name, files containing reference and target sample names, window length and step size, phasing status, number of processes, and ploidy. The model section defines the parameters of the gradient boosting quantile regressor, including the quantile level, number of estimators, tree depth, and random seed. For inference, sstar2 uses the same YAML configuration file, together with the trained model in ONNX format and the VCF file containing empirical data. Note that reference and target sample sizes as well as sequence length and phasing status used in the simulations above must match the corresponding values used for the empirical data inference.

### 2.3 Training

The train command builds the null calibration model used by sstar2. Using msprime, sstar2 simulates genomic windows under the demographic model without introgression. For each simulated window, sstar2 calculates *S*\* scores together with the count of target–reference shared variants carrying derived alleles, defined as the number of derived variants shared between the focal target genome and individuals from the reference population within the same window. These simulated data are then used to train a GradientBoostingRegressor from scikit-learn (version 1.8.0), with *S*\* scores as the response variable and the count of target–reference shared variants carrying derived alleles as the predictor. The trained model estimates *S*\* thresholds at specified null quantiles and is saved in ONNX format for downstream inference.

### 2.4 Inference

The infer command applies a trained model to empirical genotype data. For each genomic window, sstar2 calculates observed *S*\* scores and the count of target–reference shared variants carrying derived alleles. The trained quantile regression model is then used to predict the *S*\* threshold for each window at the specified null quantile. Windows with observed *S*\* scores exceeding the predicted thresholds are identified as candidate introgressed regions and written to a BED file.

### 2.5 Performance Assessment

To assess the performance of sstar2, we benchmarked it against the original implementation of sstar (version 1.2.0), QuantileRegressor from the linear_model module of scikit-learn (version 1.8.0), and RandomForestQuantileRegressor from the quantile-forest package (version 1.4.1). For sstar2, GradientBoostingRegressor was configured with quantile loss, alpha set to the specified quantile, 200 estimators, and a maximum tree depth of 3. QuantileRegressor was configured with the specified quantile, alpha set to zero, and the HiGHS solver. RandomForestQuantileRegressor was configured with 200 trees. For sstar, the ms command used an effective population size of 1,000, a reference sample size of 50 diploid individuals, a target sample size of one diploid individual, and 10,000 simulation replicates for each segregating-site count, with segregating-site counts ranging from 50 to 350 in steps of 5.

We evaluated these methods under three demographic models: ArchIE_3D19, BonoboGhost_4K19, and HumanNeanderthal_4G21 (Durvasula and Sankararaman 2019; Kuhlwilm et al. 2019; Gower et al. 2021). The focal introgression events differed across the three models (Figure 1): ArchIE_3D19 included a pulse from the source population diverging 12,000 generations ago into the target population at 2,000 generations with an admixture proportion of 0.02, BonoboGhost_4K19 included ghost-to-bonobo introgression at 20,000 generations with an admixture proportion of 0.02 from a ghost population diverging 140,000 generations ago, and HumanNeanderthal_4G21 included a pulse from Neanderthal into modern Europeans (CEU) at 1,896 generations with an admixture proportion of 0.0225, where the Neanderthal diverged 18,966 generations ago. The effective population size trajectories of populations in these models differ notably. The mutation and recombination rates were set to 1.25 × 10^−8^ per base pair per generation and 1.0 × 10^−8^ per base pair per generation for ArchIE_3D19, 1.2 × 10^−8^ per base pair per generation and 0.7 × 10^−8^ per base pair per generation for BonoboGhost_4K19, 1.29 × 10^−8^ per base pair per generation and 1.0 × 10^−8^ per base pair per generation for HumanNeanderthal_4G21, respectively.

**FIGURE 1.**
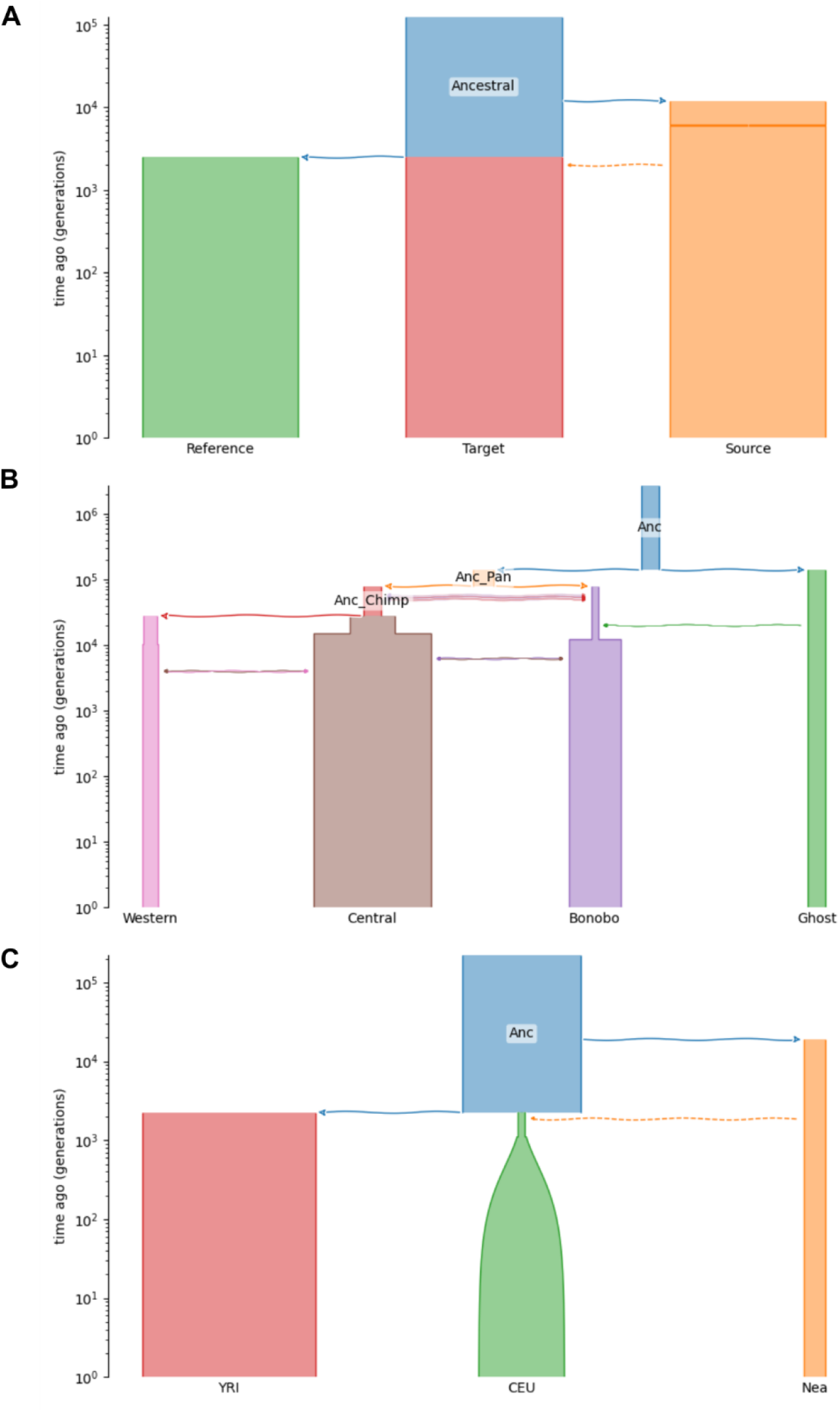
Demographic models used for performance assessment. (A) ArchIE_3D19. (B) BonoboGhost_4K19. (C) HumanNeanderthal_4G21.

For each demographic model, we used two matched versions: one without the focal introgression event for training simulations and one containing the focal event for test simulations. Training simulations were used to learn null *S*\* thresholds, whereas test simulations were used to evaluate the recovery of known introgressed tracts. Each demographic model was evaluated with 10 independent replicates, each consisting of 50 diploid reference individuals and 50 diploid target individuals. For each replicate, we generated 100,000 training windows matching the *S*\* scoring window size: 50 kb for ArchIE_3D19 and HumanNeanderthal_4G21, and 40 kb for BonoboGhost_4K19. These training windows were used to train sstar2, QuantileRegressor, and RandomForestQuantileRegressor. Test simulations were 200 Mb long to allow segment-level performance evaluation across longer genomic regions. During inference, test data were scanned using the same demographic model-specific window size as used for training, with a step size of 10 kb.

All methods were evaluated across the same set of quantile cutoffs: 0.001, 0.01, 0.1, 0.2, 0.3, 0.4, 0.5, 0.6, 0.7, 0.8, 0.9, 0.99, 0.999, 0.9999, and 0.99999. Performance was evaluated separately for each demographic model, phasing status, method, and cutoff, with results averaged across replicates. Performance was measured using segment-based precision and recall. Precision was calculated as the true positive length divided by the inferred tract length, whereas recall was calculated as the true positive length divided by the simulated introgressed tract length, where the true positive length was defined as the total overlap between inferred tracts and the known introgressed tracts extracted from msprime tree sequences.

## 3 Results

Across the simulated demographic models, sstar2 achieved the best overall performance among the evaluated methods, with the clearest improvement under the BonoboGhost_4K19 model (Figure 2). Under ArchIE_3D19, the precision–recall curves of sstar2, the original sstar, and QuantileRegressor were broadly similar for both phased and unphased data, whereas RandomForestQuantileRegressor generally showed lower performance. sstar2 showed slightly higher precision at low recall, but overall performance was similar among methods and comparable to that of other machine learning models incorporating additional input features (Huang et al. 2026), suggesting that this demographic setting is not strongly sensitive to the choice of null calibration method or machine learning model. Under HumanNeanderthal_4G21, most methods performed relatively well, with high precision maintained over a broad range of recall values, although RandomForestQuantileRegressor again performed less consistently. In this setting, sstar2 again performed similarly to the best alternative methods, indicating that replacing the original GAM-based calibration does not reduce performance in scenarios where introgressed fragments are easier to detect.

**FIGURE 2.**
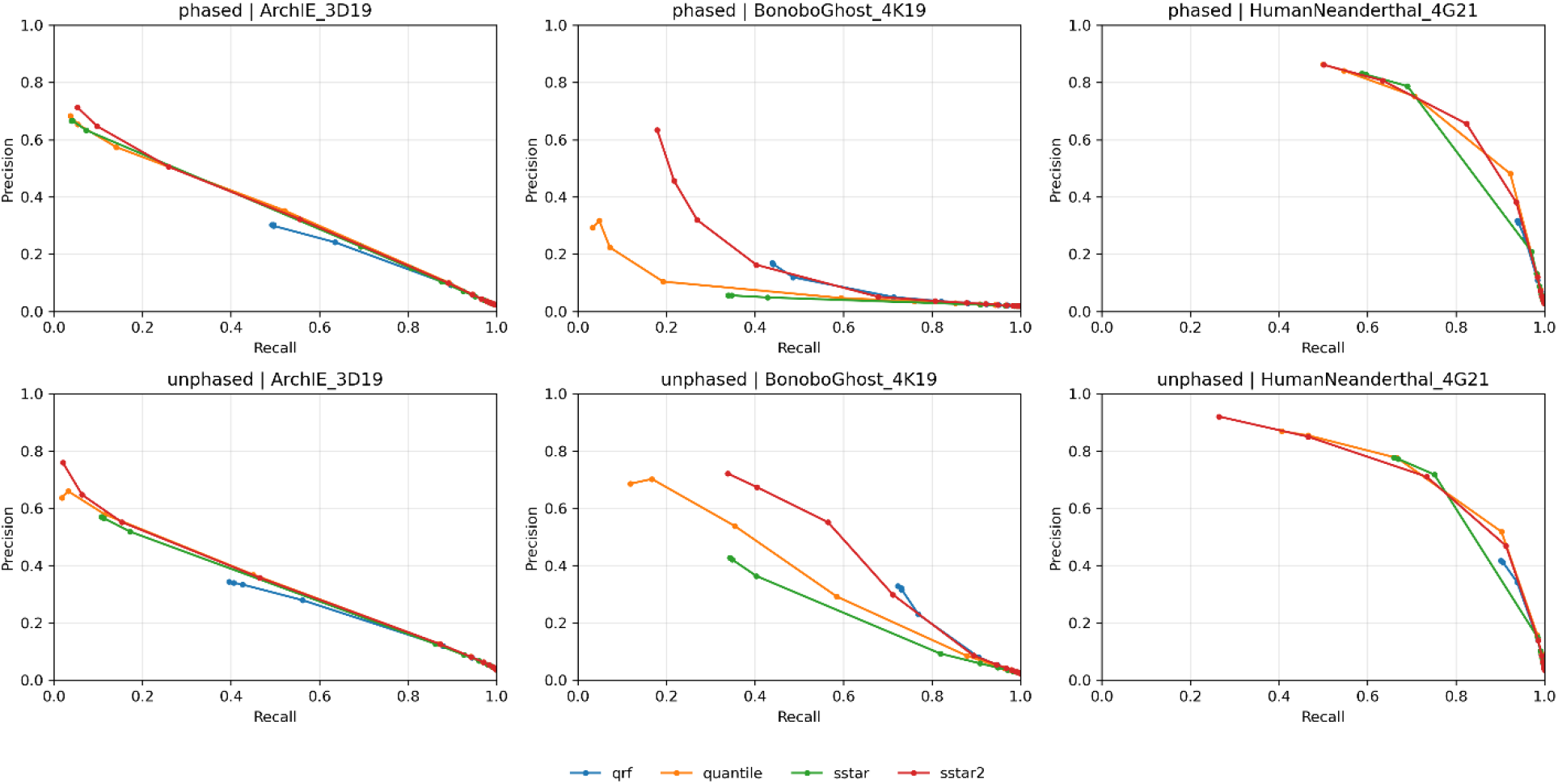
Performance comparison of sstar2 using GradientBoostingRegressor from scikit-learn, the original sstar using GAMs fitted with mgcv in R, QuantileRegressor from the scikit-learn linear_model module (quantile), and RandomForestQuantileRegressor from the quantile-forest package (qrf) across different demographic models with both phased and unphased simulated data.

The largest differences among methods were observed under BonoboGhost_4K19. In this demographic setting, which involves an ancient introgression event between deeply divergent populations, the original sstar showed low precision, especially for phased data, whereas sstar2 maintained substantially higher precision across low to intermediate recall values. The improvement was also evident for unphased data, where sstar2 achieved higher precision over a broader recall range than the original sstar and remained more stable than the alternative quantile regression approaches. These results indicate that the main advantage of sstar2 is not a uniform increase in performance across all demographic models, but improved robustness in difficult settings where the original sstar performs poorly.

Interestingly, performance was generally higher for unphased data than for phased data, particularly under BonoboGhost_4K19. This pattern likely reflects the genotype-level aggregation used in unphased *S*\* calculation. In phased data, each haplotype is evaluated separately, with each *S*\* score based on the derived variants carried by a single haplotype. In unphased data, the two haplotypes of a diploid genome are combined at the genotype level, increasing the number of variants available for *S*\* calculation. This effect may be particularly important under BonoboGhost_4K19, where the long time since introgression allows drift to substantially change the frequencies of surviving introgressed variants. Some of these variants may therefore reach high frequency or be observed as homozygous genotypes in diploid individuals, which could contribute to the higher performance of unphased genotype-level analyses.

## 4 Conclusion

Overall, sstar2 provides a modern update to the *S*\*-based framework for detecting candidate introgressed fragments from unsampled or extinct source populations. By replacing the original GAM-based calibration with quantile regression, sstar2 directly estimates *S*\* thresholds at specified null quantiles from simulated windows. Benchmarking across demographic models showed that sstar2 improves the stability of *S*\* threshold calibration in challenging scenarios, while maintaining comparable performance in easier cases. Together, these updates make *S*\*-based analyses more flexible and compatible with both phased and unphased genotype data, which will allow an easier application to empirical datasets alongside other commonly used strategies.

## Author contributions

X.H. conceived the study. A.K. and X.H. developed and evaluated the quantile regression approach. X.H., A.K., A.S., and S.C. improved sstar and implemented sstar2. X.H., A.K., and M.K. analyzed the data and wrote the manuscript.

## Acknowledgement

The authors thank the Life Science Compute Cluster at the University of Vienna and the Multi-Site Computer Austria of the Austria Scientific Computing for providing computing resources.

## Funding

M.K. was supported by the Vienna Science and Technology Fund (WWTF) [10.47379/VRG20001].

## Conflict of interest

The authors declare no conflict of interest.

## Data availability

The source code of sstar and sstar2 can be found in https://github.com/xin-huang/sstar. The Snakemake workflow for reproducing the analysis can be found in https://github.com/xin-huang/sstar-qr-analysis. All URLs were last accessed on May 31, 2026.

## Notes

### Competing Interest Statement

The authors have declared no competing interest.

